# Coupling demography and interactions reshapes the structural stability of ecological communities

**DOI:** 10.64898/2026.10.16.759606

**Authors:** Oscar Godoy, Violeta Calleja-Solanas, Fernando Soler-Toscano, José R. Portillo, Juan García-Fuentes, Daniel B. Stouffer, José A. Langa

## Abstract

The assembly of ecological communities depends on both specieś demography and the interactions among them. Ecological theory assumes that both ecological processes vary independently. However, empirical evidence demonstrates otherwise. How the structural stability of ecological communities changes when species’ intrinsic growth rates and interactions are coupled remains unknown. To address this knowledge gap, we present several seemingly disconnected examples, such as the stress-gradient hypothesis, the covariance between environment and competition, higher-order interactions, and niche packing, that actually fall along a continuum of changes ranging from positive to negative interactions. Although the specific details of each scenario differ, they all share a common feature. When demography and interactions are coupled in multi-species models of population dynamics, the limits of structural stability in ecological communities shift non-linearly, and the domain of viability increases or decreases compared to a non-coupling scenario. We mathematically demonstrate that whether these changes are positive or negative depends on how the strength of intraspecific interactions shifts relative to interspecific interactions as species’ growth rates increase. These new non-linear relationships imply two important shifts of thinking. Theoretically, the definition of structural stability must be generalized and its estimation requires the use of probabilistic measures based on ecological limits. Empirically, if abiotic and biotic barriers operate simultaneously, we need to account for such non-linear processes that can significantly modify population dynamics. Otherwise, we will not make accurate predictions of how ecological communities are assembled and biodiversity is maintained.

**Central Question:** Does the coupling between species’ demography and biotic interactions, widely observed in ecological communities, influence the opportunities for species to coexist?

**Contribution / Answer:** We apply a theoretical and modeling approach to show that the coupling modifies the structural stability of ecological communities, either expanding or reducing the opportunities for species to coexist. We derive analytical solutions to identify the conditions under which these contrasting outcomes arise for five ecological cases along a continuum from positive to negative interactions. Taken together, we provide a generalized definition of structural stability and associated methodology to study community feasibility when species interactions are not fixed.

## 1 Introduction

Ecologists understand the coexistence of multiple species within ecological communities through the concept of ecological barriers. Starting from a regional species pool, dispersal and chance represent the first barriers that species must overcome to arrive at a given site and assemble a community. Once present, they encounter the abiotic barrier, which filters species based on their ability to survive, grow, and reproduce under specific environmental conditions, such as minimum precipitation and temperature. In the last step, species must overcome the biotic barrier, which restricts their persistence at local scales if they fail to establish positive interactions with mutualistic partners or if they are unable to outcompete other species in the community. These barriers have provided a conceptual framework for understanding a wide range of ecological phenomena. They have been used to explain the limits of species distributions across continental scales (Morales-Castilla et al., 2015), the success of exotic species in becoming invasive (Blackburn et al., 2011), the emergence of alternative community diversity due to a specific sequence of events (HilleRisLambers et al., 2012), and the maintenance of species diversity within local communities (Kraft et al., 2015; Germain et al., 2018).

To test for the existence of these barriers and their consequences for the population dynamics of species, ecologists have used mathematical models in which values of each parameter inform about the relative importance of distinct processes. For instance, the abiotic barrier, commonly known as the environmental filtering metaphor, which describes the species’ inability to be established by its own, has been associated with marked reductions in demographic rates such as reproduction or survival (Kraft et al., 2015). The biotic barrier, which describes the species’ inability to establish in the presence of neighbors, has been associated with structures of species interactions that reveal the presence of natural enemies (Novotny et al., 2006), strong competitors (Chesson, 2000a), the lack of unique resources in which species can be niche differentiated (Grainger et al., 2019; Godoy et al., 2014), or the absence of positive interactions, and hence, of mutualistic partners (Godoy et al., 2026). However, the extrapolation of these ecological barriers into population models often relies on the assumption that these processes operate sequentially and largely independently (Blackburn et al., 2011; HilleRisLambers et al., 2012). In terms of modeling, this means that the variation in the values of the demographic parameters is not coupled to those of species abundances or their interactions (Fig. 1a). This assumption clashes with growing evidence that dispersal, environmental filtering, and biotic interactions are intrinsically linked (Germain et al., 2018; Matías et al., 2018). For instance, abundant species are less likely to have their spread limited by dispersal barriers (García-Callejas et al., 2021). Likewise, high abundance is typically associated with elevated reproduction and survival rates, making these species stronger competitors and less susceptible to natural enemies than species that produce little offspring (Keane et al., 2002), but see (Janzen, 1970; Connell, 1971). On the other hand, abundant species can also facilitate the establishment of other species by creating ecological niches that would not exist in their absence (Lawton et al., 1995).

**Figure 1:**
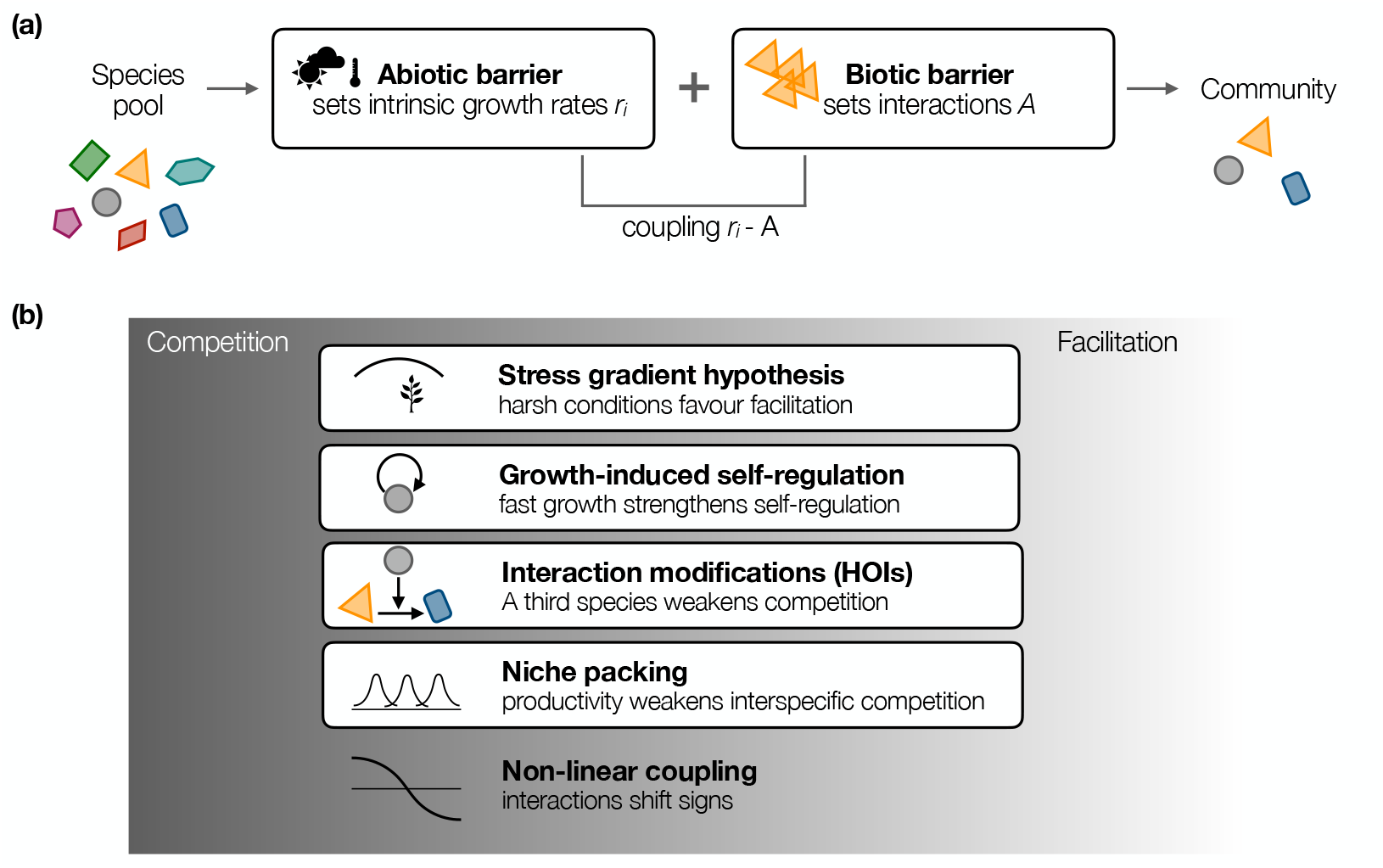
**(a)** Rather than treating abiotic and biotic barriers as independent filters, we couple them so that the strength and sign of interactions change with the growth rates the environment permits. **(b)** Four ecological processes that impose such a coupling, placed along the continuum from positive to negative interactions. In essence, these four processes are special cases of non-linear coupling of species interactions to intrinsic growth rates between interactions and intrinsic growth.

The ecological consequences of accounting for joint effects among ecological barriers remain largely unexplored, leading to an important gap in our conceptual understanding of how species coexist and communities assemble. To address that limitation when operationalizing these ideas into mathematical models, a fundamental question arises that has received little attention so far: Is the assembly of ecological communities different when model parameters are coupled than when they operate independently? Here we answer this question by exploring five types of coupling between ecological characteristics of species that emerge depending on the environmental context (Fig. 1b). These are **1) The stress-gradient hypothesis**: As environmental conditions deteriorate, species are less able to reproduce and survive (Bertness et al., 1994b; Maestre et al., 2009), and their persistence depends on the reduction of competition. It can also depend on the ability to establish positive interactions (i.e., facilitation) with other species that ameliorate local conditions and counteract the negative effects of environmental stress. **2) Covariation between environment and competition**: As formalized by the storage effect (Chesson, 2000a; Sears et al., 2007), superior competitors experience stronger intraspecific competition under favorable environmental conditions. High reproductive success in good years increases population density, which in turn strengthens self-regulation and reduces the long-term advantage of dominant species, therefore reducing the strength of the biotic barrier for other species. **3) High-order interactions**: As more species are present in a community and their abundances becomes more even, individuals of species in a community do not experience the presence of a stronger competitor due to the emergence of third species that weaken pairwise competition (Mayfield et al., 2017; Gibbs et al., 2026). **4) Niche packing reduces competition:** As productivity increases, mutualistic interactions arising from pollinators or mycorrhiza can promote species persistence by weakening the intensity of competition among plant counterparts. By increasing access to limiting resources, reducing pollen limitation, or enhancing nutrient acquisition, mutualists lessen the strength of interspecific competitive interactions (Van Der Heijden et al., 1998; Tilman, 1982; Bastolla et al., 2009). **5) Non-linear interactions**: The four previous cases describe specific mechanisms coupling demography and species interactions, but do not explicitly account for changes in interaction strength and sign. Recently, Buche et al., 2026 proposed a versatile sigmoidal function that holds the potential to unify these mechanisms along a nonlinear continuum from positive to negative interactions, where interactions change as a saturating function of productivity.

We explore these connections by coupling tools from structural stability (Saavedra et al., 2017; Godoy et al., 2018; Song et al., 2020) to community models that describe the dynamics of the species (i.e., changes in population abundances over time) and the assembly of communities (i.e., presence or absence of the species, and their relative abundance) based on two sets of parameters. These are the intrinsic ability of species to produce new individuals *r*_*i*_, and hence, to increase their abundance, and the interaction matrix *A* which contains the strength and sign of intra and interspecific interactions. We take these parameters into account because, in its simplest definition, structural stability understands community assembly and coexistence through the joint influence of species’ demography and their interactions (both sign and strength), and describes the different paths by which a community can be assembled from an empty community to its maximum levels of diversity (Letten et al., 2019; Almaraz et al., 2024; Godoy et al., 2024). With this theoretical approach, our ultimate aim is to seek a new set of general principles of community assembly that only emerge when ecological strategies among species are coupled through underlying ecological processes. Therefore, these results will provide novel insights into how the feedback between the abiotic and the biotic barriers matters for community assembly.

## 2 Theoretical framework

### 2.1 Modeling population dynamics with coupled parameters

We start with the generalized Lotka-Volterra model (LV), a well-established model that has been widely used in community ecology (Case, 2000):

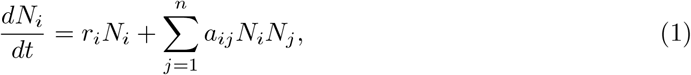

It describes the rate of change of the population size of species *i* over time 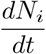 as a function of the intrinsic growth rate of species *i, r*_*i*_, and species interactions {*a*_*ij*_}. These interactions are encoded in the interaction matrix *A*, whose diagonal elements represent intraspecific interactions *a*_*ii*_ (or self-limitation) and whose off-diagonal elements are the interspecific interactions *a*_*ij*_.

Traditionally, the Lotka-Volterra model, as well as other population models such as the Ricker or Beverton-Holt models (Ricker, 1954; Beverton et al., 1957), have been used to define the dynamics of an ecological community given a particular fixed set of parameters. At each time *t*, the composition of the community is represented by the abundance column vector **N** = (*N*_1_(*t*), …, *N*_*n*_(*t*))^*T*^. If the system tends asymptotically to an equilibrium point, the final state of abundances at which the whole system stabilizes is a solution of an algebraic expression of the matrix of interactions *A* and the column vector of intrinsic growth rates **r**:

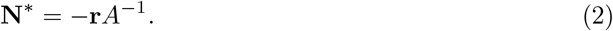

As an example, the four equilibrium points 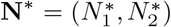 of the two-species LV model (*n* = 2 in Eq. (1)) are given by:

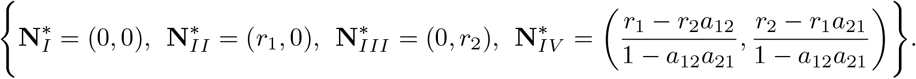

These equilibria are solely determined by the fixed species parameters. However, such equilibria will change if we allow parameters to vary.

Parameter variation emerges from biotic (same or other species densities) and abiotic (such as precipitation, nutrient, or light availability) drivers. These abiotic and biotic factors usually act simultaneously on both species demography and interactions, for example, both could be a function of precipitation *θ*: *r*_*i*_ = *f* (*θ*) and *a*_*ij*_ = *f* (*θ*). Frequently, species that in a given environment produce more offspring are also more sensitive to interspecific competition (Matías et al., 2018; Lanuza et al., 2018; Wainwright et al., 2019; Cervantes-Loreto et al., 2023), and to self-limitation (Domínguez-Garcia et al., 2026). Instead of describing this variation of parameters independently, we instead use the fact that abiotic and biotic drivers act synergistically to introduce a coupling between parameters. From this perspective, variation in one parameter, for example *r*_*i*_, directly brings about the variation of another parameter like self-limitation *a*_*ii*_, and this implies that the coupling can be described mathematically as self-limitation being a function of intrinsic growth *a*_*ii*_ = *f* (*r*_*i*_). It would also be possible to formulate coupling among parameters as an explicit temporal dependence. That is, we could explore how the variation of a parameter in one year affects the variation of another parameter in the next year by defining time-varying parameters with a certain time-lag. In mathematical terms, this approach will fall within the domain of non-autonomous systems (Chepyzhov et al., 2002; Carvalho et al., 2020; Kloeden et al., 2011; Carvalho et al., 2013). However, we do not cover such approach in this work since equilibrium points do not occur in non-autonomous systems (Godoy et al., 2024), and therefore it does not make sense to explore such scenarios with the tools from structural stability we explore here.

### 2.2 Generalizing the definition of structural stability under parameter coupling

In ecology, structural stability is mathematically defined as a measure of the set of parameters under which one can ensure that the ecosystem exhibits maximum biodiversity (Rohr et al., 2014; Saavedra et al., 2017; Godoy et al., 2018). Specifically, for the generalized Lotka-Volterra model introduced in the previous section, structural stability provides a measure for the set of intrinsic growth rate parameters that lead the system to its maximum biodiversity. However, the historical definition of structural stability no longer holds when the parameters are coupled, that is, when they are functionally dependent on one another according to an underlying ecological process. Therefore, this situation calls for a generalization of the definition of structural stability for ecologists.

When the interaction matrix is not fixed but a function of the vector of intrinsic growth rates *A* = *f* (**r**) ≡ *A*(**r**), then, for each value of intrinsic growth rates, elements of the interaction matrix change their value. The abundances **N** = (*N*_1_, …, *N*_*n*_)^*T*^ of *n* species now evolve according to the coupled LV model:

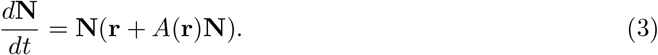

This coupling function *A*(**r**) provides the core through which we explore many of the underlying ecological processes introduced above. More generally, it refers to a broader principle that growth rates and density dependence are intrinsically coupled, as it has been previously illustrated in a wide diversity of models including the logistic growth (Stouffer, 2022) and MacArthur’s consumer–resource model (MacArthur, 1970; Chesson, 1990; Stouffer et al., 2026). Under such circumstances, the system is said to be *feasible* if there exists a steady-state solution **N**^*^ where all species persist with strictly positive abundances. By setting the time derivative to zero and assuming *N*_*i*_ ≠ 0 for all *i*, we obtain the equilibrium condition **r** + *A*(**r**)**N**^*^ = 0. Solving for the abundances yields the implicit mapping **N**^*^(**r**) = −[*A*(**r**)]^−1^**r**. Unlike in the classic model (Eq. 2), the equilibrium vector **N**^*^ now also depends non-linearly on **r** through the inversion of the varying matrix *A*(**r**).

Moreover, the set of growth rates for which all species persist is called the feasibility domain *D*_*F*_. It is formally defined as the region in the parameter space of intrinsic growth rates where the equilibrium vector remains strictly positive component-wise: *D*_*F*_ = {**r** ∈ R^*n*^ | −[*A*(**r**)]^−1^**r** *>* 0}. In the past, the feasibility domain contained vectors of intrinsic growth rates that make the system feasible. In our framework, vectors are no longer the object of study because the same vector can fall inside or outside the feasibility domain when its limits are non-linear (Fig. 2)b. Therefore, a generalized definition of the feasibility domain is the parameter space that collects every feasible point of environmental conditions, expressed through the intrinsic growth rates, under which the community retains its full diversity, and the extent of this collection of points is the quantity we want to measure.

**Figure 2:**
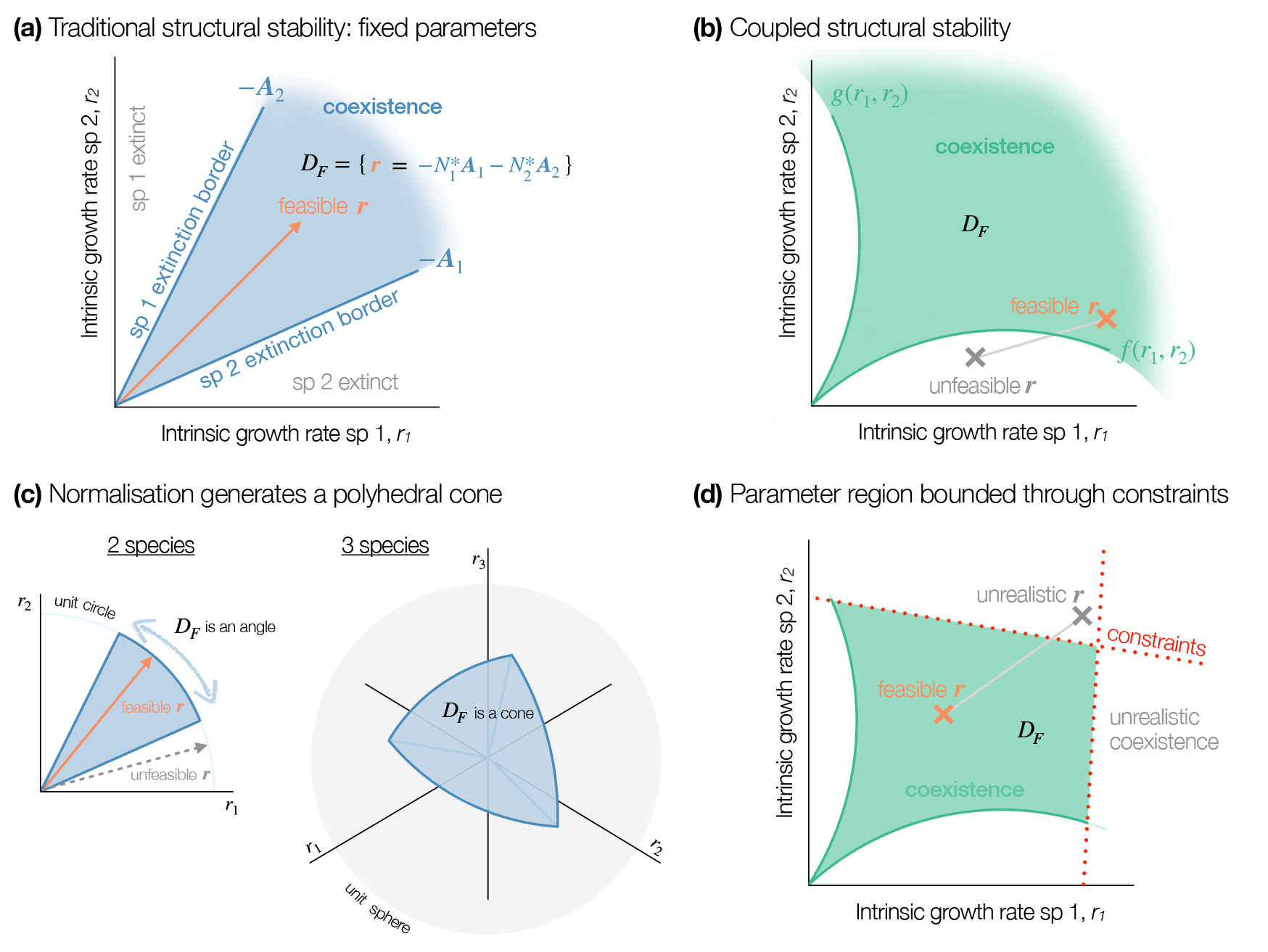
**(a)** Visualization of the feasibility domain *D*_*F*_ for a toy community of two species. The blue-brushed region represents the parameter space of intrinsic growth rates, generated by the negative column vectors (spanning vectors) of the interaction matrix **A**_1_ and **A**_2_, for which species coexist. The orange arrow is a hypothetical vector of intrinsic growth rates inside *D*_*F*_. When the intrinsic growth rates of species cross the extinction borders, the equilibrium point is not feasible since some species go extinct. **(b)** When matrix elements are coupled with intrinsic growth rates, the extinction borders bend. Given a collection of points containing information of species’ intrinsic growth rates, the community can be unfeasible (gray point) or feasible (orange point) depending on where they sit, even if along the same line. **(c)** The light blue unit circle and unit sphere correspond to the normalized parameter space for 2 and 3 species, respectively. For any number of species, the normalized solid angle of the cone (simply an angle in 2-species systems) of the feasible domain is the fraction of the area in the unit hypersphere occupied by the cone (blue-brushed region), which is the probability of sampling uniformly a vector of intrinsic growth rates **r** on the unit hypersphere inside the feasibility domain. A hypersphere for 2 dimensions (species) is a circle, and a sphere for 3 species. **(d)** Even though the projection onto a hypersphere makes no sense when parameters are coupled, we can still measure the size of the *D*_*F*_, taking into account that, depending on the context, the feasibility domain may be constrained by more than just the interaction matrix. For example, ecological constraints imposed by realistic intrinsic growth rates (dotted red lines) and projected prey and predator abundances may define a collection of points that fall with in curved boundaries of the *D*_*F*_ (green curves). Hence, the collection of points that fall outside *D*_*F*_ can either create an unfeasible scenario or unrealistic based on the imposed ecological constraints.

To see why coupling changes *D*_*F*_ ‘s shape and size, consider first the classical case of a constant matrix. The key point is that for the requirement of each species having 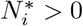, we have a linear inequality that divides the parameter space in two along a flat boundary through the origin. The feasibility domain is what remains once all requirements are imposed at once **N**^*^ = −*A*^−1^**r** *>* 0: a polyhedral cone with flat faces (boundaries), one per species, meeting at the origin (Fig. 2a). Crossing a boundary means that one species reaches zero abundance and goes extinct. When parameters are instead coupled, such as when the matrix depends on growth rates, *A*(**r**), each of those requirements still marks the extinction of one species, but it is no longer linear, because the coefficients that define it now change as **r** moves. The faces of the cone bend into curved surfaces whose exact shape follows from the functional form of the coupling (Fig. 2b). The boundaries can bend outward, admitting growth rates that would be unfeasible under a constant fixed matrix, or inward, excluding ones that would have been feasible. As the coupling change the domain to no longer be a cone, the geometric shortcuts built on that assumption no longer apply.

### 2.3 Solving the limitation of traditional geometric measures to explore structural stability when ecological strategies are coupled

The traditional geometric approach for quantifying structural stability relies on the demonstration that the feasibility domain *D*_*F*_ forms a polyhedral cone rooted at the origin. In systems with a constant interaction matrix *A*, the feasibility condition is scale-invariant with respect to the growth rates **r**. That is, if a vector **r** leads to a feasible community, any scaled vector *cr* (for *c >* 0) is also feasible. This conical symmetry allows the problem to be simplified to measuring a finite surface area as the *D*_*F*_ size, which is the intersection of the cone with the unit hypersphere (Saavedra et al., 2017; Song et al., 2020). The size of the *D*_*F*_ becomes the area of this intersection, which is the solid angle between the column vectors of the interaction matrix for a two-species system (Fig. 2c). However, when the interaction matrix depends on the growth rates, *A*(**r**), this scale invariance is broken. The feasibility boundaries become irregular hypersurfaces, and the domain is no longer a cone. Consequently, the intersection of this domain with a unit hypersphere becomes an arbitrary cross-section that fails to capture the true shape or “size” of the feasible region. An **r** vector might be feasible at a small magnitude but unfeasible at a larger magnitude due to the shifting interactions, rendering the spherical projection meaningless as a global measure of stability (Fig. 2d).

Furthermore, we face a computational limitation. Calculating the exact volume of a high-dimensional region bounded by non-linear, implicitly defined hypersurfaces is analytically intractable for all but the simplest functional forms of *A*(**r**). Standard integration techniques require explicit boundary definitions, which are often unavailable when the boundaries are roots of complex nonlinear equations. To address this limitation, the solution we present here is a numerical approach that can handle arbitrary constraints without relying on the geometric simplifications of linearity.

### 2.4 A bounded Monte Carlo approach to estimate structural stability

To overcome these topological and analytical hurdles, we adopt a probabilistic measure of structural stability within ecologically relevant constraints (Song et al., 2018). This probabilistic measure of structural stability has been considered before when accounting for the effect of environmental variability and multitrophic interactions on defining the feasibility domain (Song et al., 2020; Saavedra et al., 2020; Cervantes-Loreto et al., 2023; Buche et al., 2024). Instead of measuring an abstract solid angle, we define a bounded parameter region for the intrinsic growth rates, 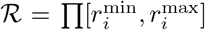, based on empirical ecological constraints, for instance, maximum reproduction rates that are physiologically and metabolically possible. The structural stability can be quantified as the proportion of a reference region of intrinsic growth-rate space in which all species have positive equilibrium abundances (Fig. 2d). Let *S* denote the number of species and **r** = (*r*_1_, …, *r*_*S*_)^⊤^ the vector of intrinsic growth rates. We consider a reference region *ℛ* ⊆ ℝ ^*S*^ with finite, nonzero volume *V* _*ℛ*_ = Vol(*ℛ*). For a given growth-rate vector **r** ∈ *ℛ*, the equilibrium abundance vector is given by

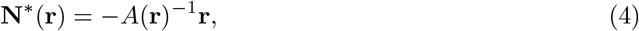

assuming that the interaction matrix *A*(**r**) is invertible. Here, *A*(**r**) is an *S* × *S* interaction matrix whose element *a*_*ij*_(**r**) describes the effect of species *j* on the per-capita growth rate of species *i*.

The feasible domain within *ℛ* is defined as

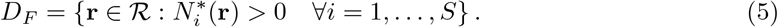

The relative structural stability is then expressed mathematically as

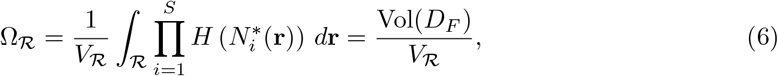

where *H*(·) is the Heaviside function, defined as

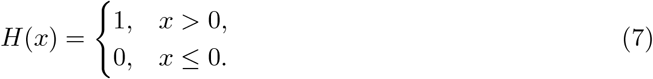

This convention ensures that an equilibrium abundance equal to zero is considered infeasible. Equivalently, the product of Heaviside functions can be expressed as the indicator function of the feasible domain:

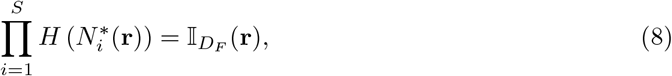

where

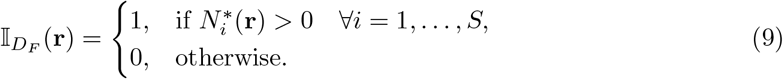

Thus, integrating the indicator function over *ℛ* yields the volume of the feasible domain within the reference region:

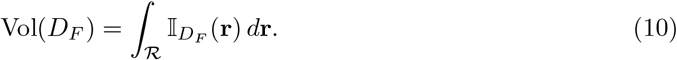

Rather than solve the integral analytically, we estimate this volume using Monte Carlo integration. Specifically, we generate *K* growth-rate vectors independently and uniformly distributed over *ℛ*, and evaluate the feasibility condition following the indicator function 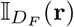. Let *N*_*F*_ denote the number of sampled points for which all equilibrium abundances are strictly positive. The absolute volume of the feasible domain is then estimated as

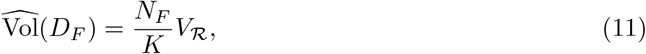

whereas the corresponding relative structural stability is estimated as

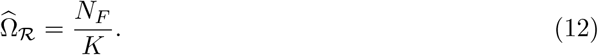

To illustrate our approach, we define coupling functions that are contiguous and ecological constraints that only limit the boundaries of the *D*_*F*_ (Fig. 2). However, these domains can potentially contain discontinuities based on how the coupling functions are defined and the ecological constraints imposed (Appendix). These holes can vary in grain (small versus big), complexity (regular versus irregular), and position (inside versus along the boundary of the domain). The probabilistic measure we adopt here is flexible enough to accommodate all such irregularities without requiring their analytical characterization. The estimate of absolute feasible volume depends on the extent and geometry of the reference region *ℛ*, which must therefore be specified when comparing systems.

## 3 Results on coupling species’ growth rates and interactions across ecological processes

We begin by exploring two well-known cases in the literature of two-species systems whose ecological strategies can be modeled using coupling parameters, and illustrate how the species’ opportunities for coexistence change under this new perspective, section 3.1. In both cases, we have selected situations known to promote species coexistence, either because species are buffered from extinction by facilitation or because species limit their dominance and therefore do not exclude inferior competitors through self-limitation. Therefore, we allow interaction coefficients to vary in sign (positive versus negative) and strength (strong versus weak). Although they are commonly treated in the literature as independent phenomena, they are tightly connected in nature, and we will present in the subsequent sections 3.2-3.3 two cases with more than two species indicating how we can model a continuum from facilitative to competitive interactions (Buche et al., 2025b).

### 3.1 Stress gradient hypothesis and growth-induced self-regulation: Two sides of the facilitation-competition continuum

The first case is the stress gradient hypothesis (SGH) that posits a monotonic increase in the role of facilitation with increasing environmental harshness (Bertness et al., 1994a; Maestre et al., 2009). Such severity and the prevalence of facilitation have been observed across many different plant ecosystems. Indeed, some ecosystems include stressful environments such as drylands, in which nurse plants facilitate seedling establishment and survival by ameliorating water and radiation stress (Berdugo et al., 2019; Calleja-Solanas et al., 2025; Caligiuri et al., 2025), or cold environments (alpine and forest at high latitudes) in which cushion plants facilitate the survival of neighbors by protecting them from herbivory and freezing temperatures (Choler et al., 2001; Godoy et al., 2015). Yet, we would like to stress that facilitation is also prevalent in other ecosystems that do not experience such stressful conditions (Buche et al., 2026).

In mathematical terms, we model the stress gradient hypothesis as follows: the intrinsic growth rates are the proxy for the harshness of environmental conditions; thus, as conditions deteriorate (*r*_*i*_ ↓), the off-diagonal of the interaction matrix containing the per-capita interspecific interactions *a*_*ij*_ changes towards positive values (prevalence of facilitation). Therefore, diversity can be maintained because facilitation offsets the reduction of intrinsic growth rates arising from environmental stress. In other words, facilitation is a mechanism to avoid species’ environmental filtering. Using the Lotka-Volterra model, the off-diagonal elements of the interaction matrix are coupled with the intrinsic growth rates in such a way that *A* is defined as:

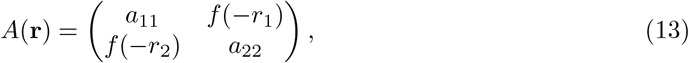

where *a*_*ii*_ *<* 0 and *f* (*x*) is a monotonically increasing function. Consequently, *f* (−*r*_*i*_) increases as *r*_*i*_ becomes more negative, representing an increase in facilitation with increasing abiotic stress. According to this formulation, we observe that as facilitation increases with abiotic stress, the domain expands into regions where *r*_*i*_ *<* 0, Fig. 3a (see Appendix for how the boundaries were computed). This finding reflects mutualistic rescue, and it is consistent with previous modeling of plant-mutualistic systems and horizontal plant communities (Rohr et al., 2014; Godoy et al., 2026).

**Figure 3:**
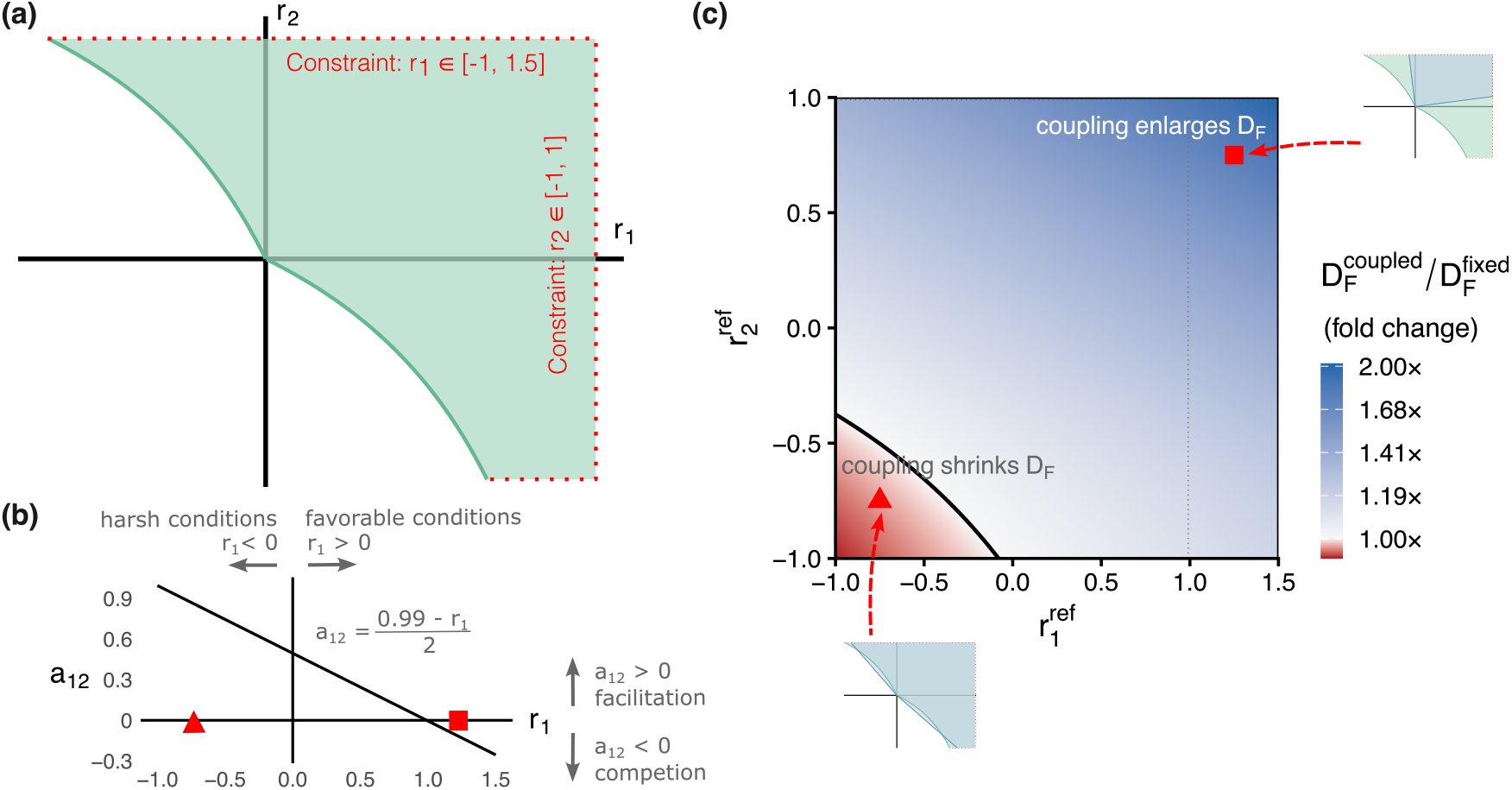
Stress Gradient Hypothesis. **(a)** Coupled feasibility domain 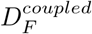 intersected with constraints on the intrinsic growth rates given by the regions *r*_1_ ∈ [−1, 1.5] and *r*_2_ ∈ [−1, 1]. This single domain is the coupled reference against which every constant matrix in (c) is compared. **(b)** Functional form of the coupling between intrinsic growth rates and interspecific interactions. The triangle and square mark the two reference conditions highlighted in (c). **(c)** Comparison of the coupled system with the family of uncoupled models it contains. For each reference condition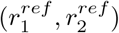, the interspecific coefficients are frozen at the values the coupling assigns there. The size 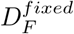 of the resulting constant matrix is computed over the same constraints. Color shows the ratio 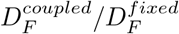 as a fold change on a log scale, and the solid black line is the contour along which the two coincide. The gray dotted line marks 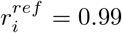, where the frozen coefficients change sign. Note that the axes identify which constant matrix serves as the fixed reference, not the growth rates of a community: the coupled domain is one number, and it is the same number everywhere in the panel. Insets are the feasibility domains for the two marks (**r**_Δ_ = (−0.75, −0.75) and **r**_□_ = (1.25, 0.75)) in blue overlapping 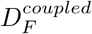 (in green).

Importantly, unlike previous models where fixed interactions yield straight boundaries (Fig. 2a), coupling the interaction coefficients to the intrinsic growth rates through *f* (−*r*_*i*_) generates curved boundaries of the feasibility domain bending outward (Fig. 3a). This means that a linear increase in facilitation as conditions deteriorate (Fig. 3b) produces a non-linear expansion of the range of intrinsic growth rates *r*_*i*_ compatible with the persistence of both species.

Quantifying that enlargement requires care, because the coupled and the fixed cases are not directly comparable one-to-one. The coupled system of Eq. (13) has a single interaction matrix *A*(*r*) and therefore a feasibility domain whose size within some constraints is a single number 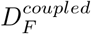. How to mathematically formalize the linear dependencies between species’ performance and facilitation hypothesized by the SGH remains unclear, as no prior work has explicitly addressed this relationship. For illustration purposes, we choose the coupling of 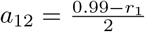 and 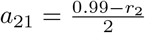 with *a*_11_ = *a*_22_ = −1 (Fig. 3b), but we acknowledge that this decision is arbitrary and further coupling should be done based on empirical information. Under this parameterization, the coupled feasibility domain is 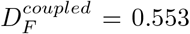 of the bounded parameter region (the green region of Fig. 3a). An uncoupled model, by contrast, requires us to commit to one constant fixed matrix, and the coupling supplies a different one for every intrinsic growth rate point **r**^*ref*^. We therefore compare the coupled system against the whole family of uncoupled models it contains. For each reference condition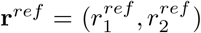, we freeze the interspecific coefficients at the values the coupling assigns there, and we estimate the size 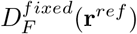 of the resulting domain with a constant matrix over the same bounded region. Ecologically, each member of this family represents a measurement of species interactions under one set of conditions, which is assumed to hold for every other condition. Fig. 3c maps the ratio 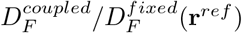 across all possible reference points given our constraints.

Coupling enlarges the feasibility domain for 93% of the reference conditions we examined, by up to a factor of 2.02 when interactions are fixed at the most favorable corner of the parameter region (the square in Fig. 3b and c gives a 1.76-fold enlargement). The exception is a narrow corner in which the reference growth rates are strongly negative, where the fixed matrix is already facilitative, and it slightly outperforms the coupled system (the triangle shape in Fig. 3b and c gives a 0.97-fold shrink). The value 0.99 keeps det*A*(**r**) ≥ 0.01 *>* 0 throughout, so a unique equilibrium exists everywhere in the bounded region. Overall, an observer who estimates interactions at a single environmental condition (**r**) and assumes them fixed will in general underestimate the opportunities for these two species to coexist.

In contrast to facilitation, competition can also promote diversity, as documented by a large body of theoretical and empirical work. A well-known example is the positive covariance between fitness and self-regulation, in which competition can be represented as a function of species’ intrinsic growth rates. This covariance describes how strong competitors do not disproportionately benefit from favorable conditions and therefore are prevented from dominating local communities. In particular, a species that performs well in a given site (i.e., high *r*_*i*_) does not obtain as much benefit as expected in terms of population growth rates, because the strength of self-limitation also increases (i.e., high *a*_*ii*_) (Chesson, 2000b; Snyder et al., 2004). This covariance can underlie physiological trade-offs where rapid growth leads to inefficient resource use, waste accumulation, or physical crowding. For instance, in microbial systems this phenomenon mirrors the so-called rate-yield tradeoff, in which fast-growing organisms such as fermenting yeast often utilize resources inefficiently, hitting carrying capacities imposed by their own toxic byproducts or nutrient depletion more quickly than they can exclude more efficient but slower competitors (Frank, 2010; Wortel et al., 2018).

In ecology, this covariance is a key element of the storage effect that is attributed to maintaining diversity in spatially variable environments because poor performance in some spatial locations is ameliorated by better performance elsewhere (Chesson, 2000a; Sears et al., 2007). For illustrative purposes, to analyze a system in which there is a positive covariance between intrinsic growth rate and self-regulation, the positive covariance between intrinsic growth rate and self-regulation can be defined as:

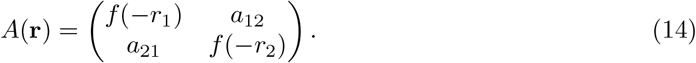

where the diagonal terms *a*_*ii*_ of the competition matrix depend quadratically on *r*_*i*_ (Fig 4a). The quadratic form is chosen because it provides the simplest nonlinear function that allows the strength of self-regulation to vary with the intrinsic growth rate while remaining negative over the range of *r*_*i*_ considered, and fits the empirical observations that as intrinsic growth rates increase the magnitude of intraspecific competition *a*_*ii*_ rises sharply (Matías et al., 2018; Wainwright et al., 2019; Van Dyke et al., 2022). Importantly, the quadratic form is not intended to represent a unique mechanistic relationship between environmental conditions and competition, but rather to provide a simple example of how such coupling can generate nonlinear changes in the feasibility domain. Note that this relationship also emerges with other functions in which the *a*_*ii*_ terms do not depend quadratically on the intrinsic growth rates.

**Figure 4:**
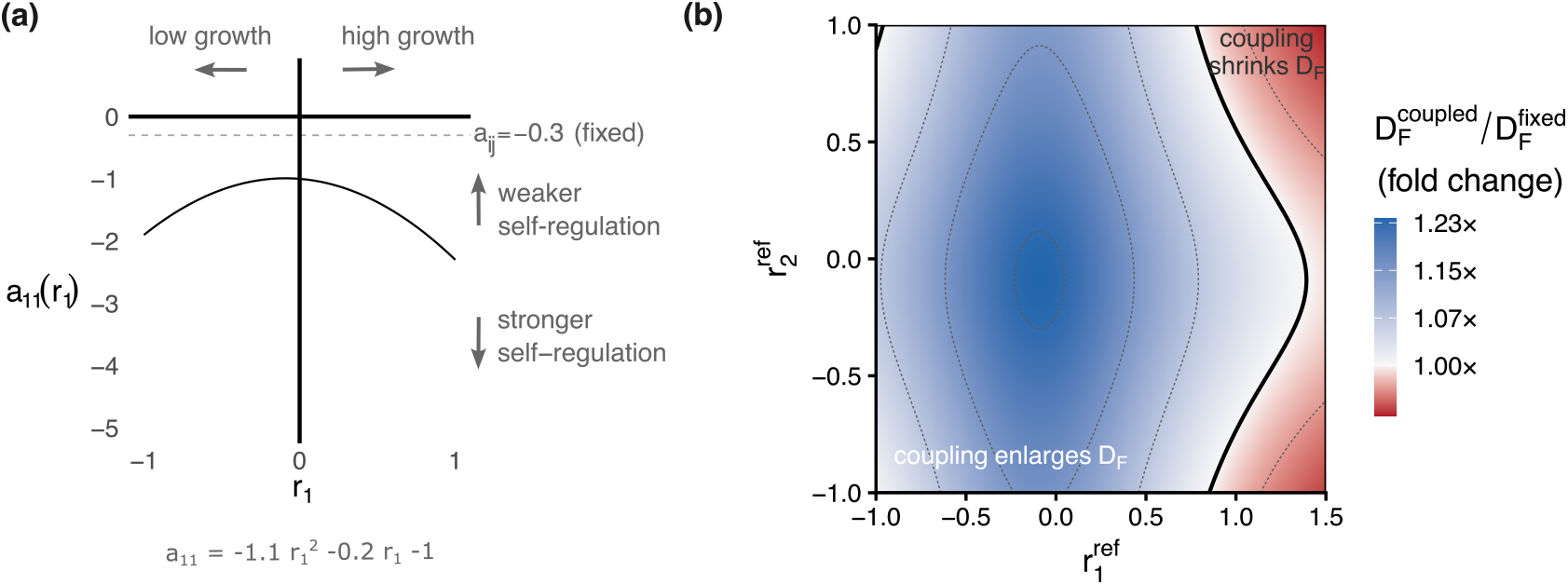
Growth-induced self-regulation. **(a)** Functional form of the coupling between intrinsic growth rates and self-regulation 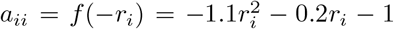. **(b)** Comparison of *D*_*F*_ of the system in Eq. 14 and its fixed versions for different values of **r**^*ref*^. The intrinsic growth rates were bounded within the same ranges as in Fig. 3b, *r*_1_ ∈ [−1, 1.5] and *r*_2_ ∈ [−1, 1], and interspecific interactions were set to *a*_12_ = *a*_21_ = −0.3.

The main result we observe here is very similar to the role of facilitation in promoting species coexistence. That is, the feasibility domain expands outward, creating a region much broader than the standard cone when intrinsic growth rates and self-limitation are not coupled (Fig. 4a). Applying the same comparison as for the stress gradient hypothesis, we contrast the coupled system of Eq. 6, whose domain occupies 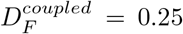 of the bounded parameter region, against the family of uncoupled fixed models obtained by freezing the matrix at each reference condition. Coupling enlarges the feasibility domain for about 84% of these reference conditions, by up to 1.24-fold (Fig. 4b). This relaxation of the feasibility constraint occurs because self-limitation (|*a*_*ii*_|) strengthens with growth much faster than the interspecific competition, which we hold constant in this case (|*a*_*ij*_| = 0.3). Consequently, the system becomes highly structurally stable because the matrix of species interactions becomes diagonally dominant at high growth rates **r**.

### 3.2 Species coexistence via high-order interactions

For decades, ecologists have largely considered pairwise interactions between species as the fundamental unit of analysis for understanding community dynamics and species coexistence (Fig. 5a). However, accumulation of theoretical and empirical evidence suggests that the outcome of an interaction between two species can be modified by the presence or abundance of a third species, giving rise to higher-order interactions (HOIs) in the sense of interaction modifications (Billick et al., 1994; Wootton, 1994a; Levine et al., 2017). HOIs can alter both the strength and even the sign of pairwise interactions, thereby influencing community dynamics in ways that cannot be captured by traditional pairwise models (Levine et al., 2017; Mayfield et al., 2017). In pairwise models, interactions are typically represented as the per-capita effect of one species on the growth of another, such as (1*/N*_2_)*dN*_2_*/dt* = *f* (*N*_1_), and, in the traditional LV model, (1*/N*_2_)*dN*_2_*/dt* = *a*_21_*N*_1_. The ecological importance of HOIs has received increasing attention in recent years, with works documenting their occurrence in both experimental and natural communities (Mayfield et al., 2017; Letten et al., 2019; Buche et al., 2024), but see Calleja-Solanas et al., 2026 and Terry et al., 2017. However, most of the current theoretical work considers that the strength of HOIs depends on the abundance of a third species. Here, we consider a complementary form of HOI in which the interaction between two species depends solely on the presence or identity of a third species, rather than on its abundance. Such dependence can arise when a third species modifies the phenotype, behavior, resource environment, or abiotic conditions experienced by the interacting pair. Mathematically, this can be represented as (1*/N*_2_)*dN*_2_*/dt* = *f* (*N*_1_, *N*_3_), for example, (1*/N*_2_)*dN*_2_*/dt* = *b*_213_*N*_1_*N*_3_ when the effect scales with the abundance of the third species. Alternatively, if the effect is triggered simply by the presence of species 3, the interaction coefficient can be written as (1*/N*_2_)*dN*_2_*/dt* = *δ*_3_*a*_21_*N*_1_, where *δ*_3_ = 1 when species 3 is present and *δ*_3_ = 0 otherwise. An equivalent formulation is to couple the interaction coefficient to the intrinsic growth rate of the third species, (1*/N*_2_)*dN*_2_*/dt* = *f* (*r*_3_, *N*_1_), thereby allowing species growth to determine the interaction structure (Fig. 5b).

**Figure 5:**
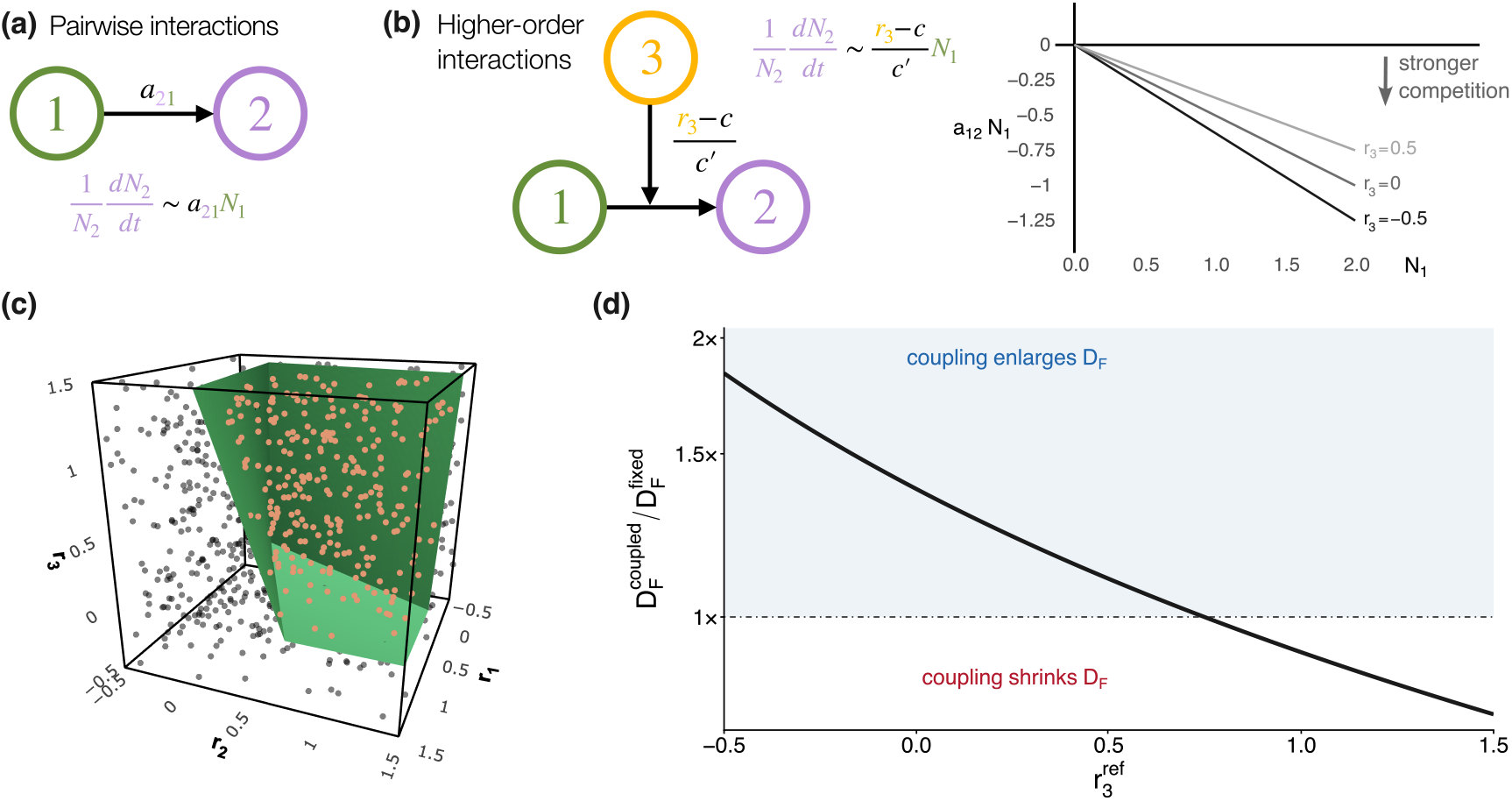
Coexistence mediated through interaction modifications with HOIs. **(a)** In a pairwise scenario, species 1 affects the per capita growth rate of species 2 through a direct effect (*a*_21_). **(b)** In a higher-order scenario, species 3 modifies the effect of species 1 on species 2, with a strength that depends on the growth rate of the initiator species (species 3). **(c)** The intersections of the green planes of constraints *r*_*i*_ ∈ [−0.5, 1], represent the available values of intrinsic growth rates. Orange (gray) dots denote points inside (outside) the feasibility domain estimated by the Monte Carlo algorithm with 1000 points just for illustrative purposes. The species have −1 as self-regulation and *a*_12_ = *a*_21_ = 0.5 − *r*_3_*/*4 and zero elsewhere. The estimations of *D*_*F*_ sizes had 10^7^ points throughout the work. **(d)** Comparison of the coupled system with the family of uncoupled models it contains. The curve shows the ratio 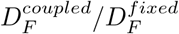 as a fold change on log scale; the dash-dotted line at 1 marks equality between the coupled and uncoupled cases, and blue shading indicates the reference conditions for which coupling enlarges the feasibility domain, following the same convention as Figs. 3c and 4b.

To explore this coupling, we define the interaction modification as *f* (*r*_3_, *N*_1_) = *N*_1_(*r*_3_ − *c*)*/c*^′^, where *c* and *c*^′^ are positive constants, so that the structure of a matrix of interactions for three species is as follows:

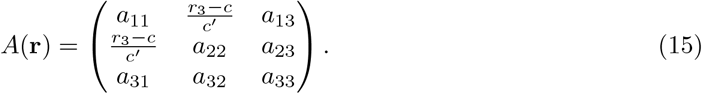

In this formulation example, HOIs emerge from the presence of a third species (the initiator) which changes the per capita effect of a competing species 1 (the transmitter) on a focal species 2 (the receiver) (following nomenclature of Buche et al., 2024).

The presence of this HOI means that while species 1 and 2 are competitors in isolation (values of *a*_12_ = *a*_21_ = −*c* at *r*_3_ = 0 in Eq. (15)), the strength of such competition is reduced as species 3 gradually thrives (*r*_3_ ↑, Fig. 5b). Many ecological examples drive this three-species process (Wootton, 1994b). For instance, mycorrhizal fungi act as an initiator of the HOI, redistributing resources between plant species, effectively lowering competition for resources and reducing asymmetries in competitive ability (Willing et al., 2024). Another example comes from cross-feeding in microbial systems in which a bacteria species breaks down complex polysaccharides into abundant simple sugars, relieving the scramble competition between two specialized consumers that would otherwise be involved in competitive exclusion (Culp et al., 2023). A final example of HOIs that does not involve an increase in resources is when a top predator initiates a change in the behavior of two prey species that compete for common resources to graze (Werner et al., 2003).

Under this scenario, we show that HOIs reshape the feasibility domain. As the intrinsic growth rate of the HOI initiator increases (species 3), the competition between species 1 and 2 weakens, and the domain expands to such an extent that it creates a buffer zone in which the two competing species can coexist over a broader range of their own growth rates (Fig. 5c). Higher-order interactions can promote species coexistence and expand the feasibility domain nonlinearly (Fig. 5d). Applying the same comparison as in the previous examples, we compare the coexistence opportunities in the coupled system against the family of uncoupled models. Coupling enlarges the feasibility domain for reference conditions below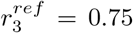, by up to 1.83-fold, and reduces it above that value, by down to 0.79-fold Fig. 5d. These results indicate that the assessment of the importance of HOIs depends on the community with uncoupled parameters used as the benchmark. If the growth rate of the initiator is high, we would infer weaker competition and hence a larger feasibility domain, thereby underestimating the contribution of the HOI. Meanwhile, if the growth rate of the initiator is low, we would infer stronger competition and a smaller feasibility domain, thereby attributing a larger relative contribution to the HOI. Thus, the estimated importance of the HOI depends on the conditions under which interactions are measured, echoing the same lesson from the stress-gradient and self-regulation examples and reinforcing the point that a single fixed interaction matrix is not a neutral point of comparison.

### 3.3 Species coexistence via niche packing

Along productivity gradients, species can vary their interactions (Rees, 2013). When resources are scarce, species tend to greatly overlap in their niches, which intensifies the prevalence of competition among species. As productivity increases, species frequently partition resources along spatial, temporal, or functional axes, reducing competition between species but increasing within-species competition, although the precise relationship depends on the mechanisms underlying the productivity gradient. There is a plethora of examples documenting this process for annual plant communities (Matías et al., 2018), tropical rain forests (Fort et al., 2013), and insect communities (Novotny et al., 2006).

If we assume that the amount of total biomass depends of the number of individuals and the ability of the species to grow their populations within communities (*r*_*i*_) as commonly observed in forest and grassland ecosystems (Hector et al., 2002; Storch et al., 2018), we can examine a scenario in which interspecific competition is inversely related to the overall system productivity 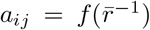, where 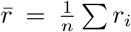. The basic underlying mechanism of this scenario is to assume that as the productivity of the system increases, the denominator grows, effectively reducing the relative strength of competition between species (|*a*_*ij*_| →0) compared to within species (Fig. 6a,b). This captures the hypothesis that high-productivity environments support more complex habitat structures or resource diversity, allowing species to partition niches more effectively and avoid direct conflict (Wright, 2007).

**Figure 6:**
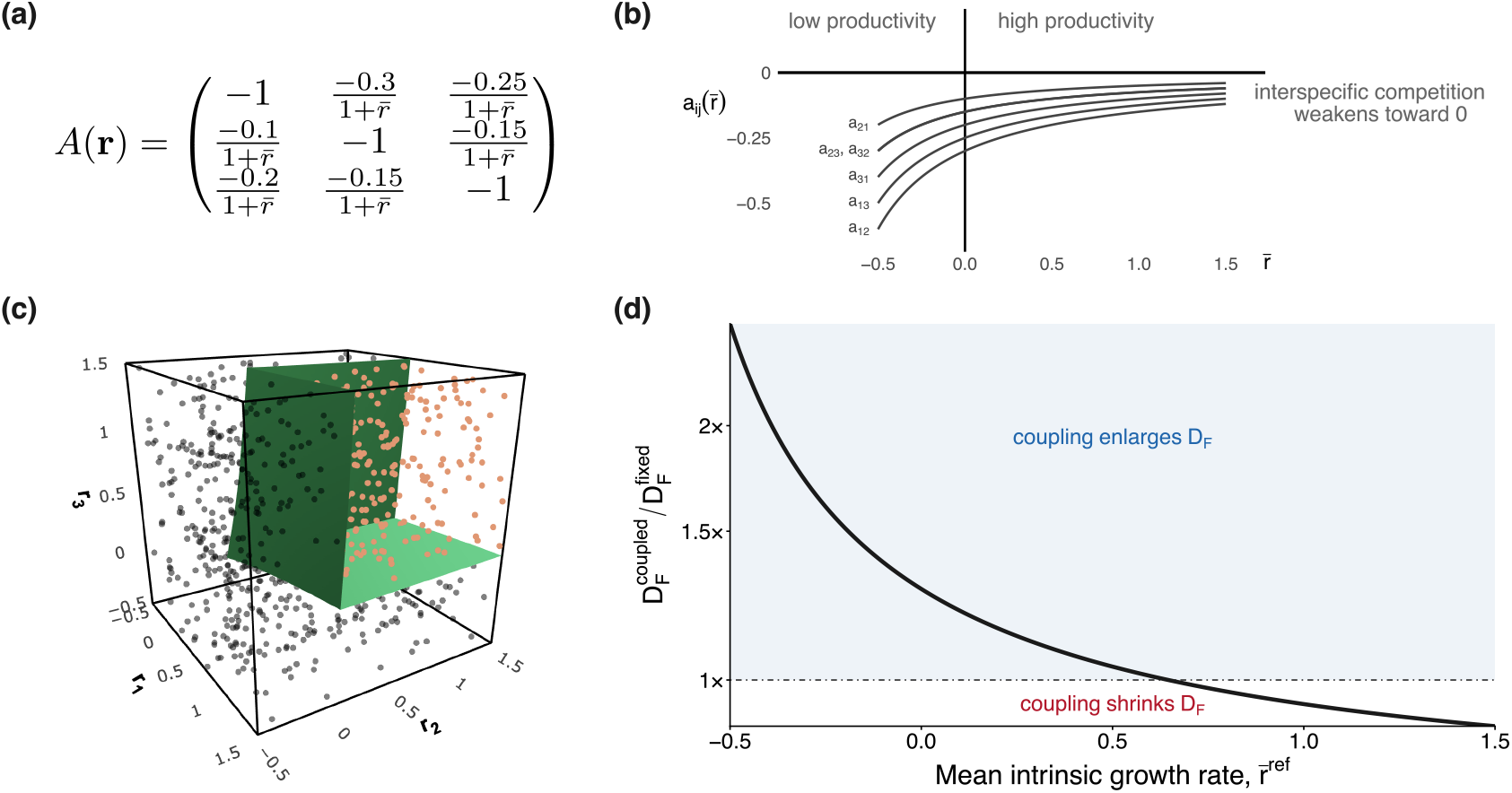
Coexistence via niche packing. In this example, the interaction matrix follows the expression in panel **(a)**, in which all interspecific terms are scaled so that competition between species weakens as ecosystem productivity rises, while self-limitation is held at *a*_*ii*_ = −1. Intrinsic growth rates are constrained to *r*_*i*_ ∈ [−0.5, 1.5] for all three species. **(b)** Functional form of the coupling between overall productivity and interspecific interactions. **(c)** Feasibility domain of the coupled system intersected with these constraints, whose green planes delimit the available values of intrinsic growth rates. Orange (black) dots denote points inside (outside) the domain, estimated by the Monte Carlo algorithm. Its volume is about 29% of the bounded parameter region, and it is the coupled reference against which every constant matrix in panel (b) is compared.**(d)** Comparison of the coupled system with the family of uncoupled models. For each reference mean productivity 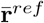, the interspecific terms are frozen at the value the coupling assigns there. The curve shows the ratio 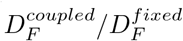 as a fold change on log scale; the dash-dotted line at 1 marks equality between the coupled and uncoupled cases; and blue indicates the reference conditions for which coupling enlarges the feasibility domain. The horizontal axis identifies which constant matrix serves as the benchmark, not the mean growth rate of a community. Note that 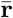 denotes the mean growth rate of a community, whereas 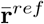 only labels which constant matrix serves as the benchmark; each frozen matrix is evaluated over the full constraint region regardless of the value used to define it.

Similar to the findings of the previous cases, we find that this coupling of the productivity of the ecosystem and niche partitioning leads to an expansion of the feasibility domain (Fig. 6c). While conditions of low growth enforce strict competitive exclusion boundaries, the coupling promotes larger coexistence opportunities than the fixed matrices. As in the previous case of higher-order interactions, the declining 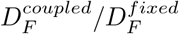 ratio reflects an improvement in the reference fixed matrix rather than a weakening of niche packing (Fig. 6d). That is, an observer who measures competition in a productive year will infer weak interspecific effects that generate a 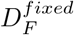 larger and more similar in size to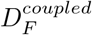. Together with the previous examples, this establishes a consistent pattern across all four processes. Coupling relaxes the constraints on diversity relative to most fixed matrices one could estimate.

### 3.4 A common currency for modeling coupling between species’ growth rates and interactions across ecological strategies

Although these four scenarios can be placed along a continuum from positive to negative interactions, each has been formalized with a different functional form. In this final step, we move beyond these specific formulations and introduce a common mathematical representation (sigmoidal function) of parameter coupling (Buche et al., 2025b; Buche et al., 2026). Here, the asymptotic bounds of the sigmoid serve as scaling limits rather than biologically prescribed values, allowing the strength and direction of coupling to vary continuously between facilitation and competition. In natural systems, environmental gradients and overall productivity often induce smooth, threshold-like transitions in species interactions that cannot increase forever, rather than unbounded linear shifts. To capture this behavior, we define the strength of interspecific interactions, 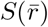, as a sigmoidal function of the community’s mean intrinsic growth rate:

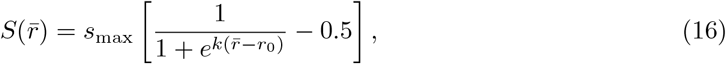

where *s*_max_ controls the amplitude of the transition, *k* its steepness, and *r*_0_ the value of 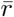 at which interactions switch sign. In Fig. 7a, we use *s*_max_ = 1, *k* = 4, and *r*_0_ = 0:

**Figure 7:**
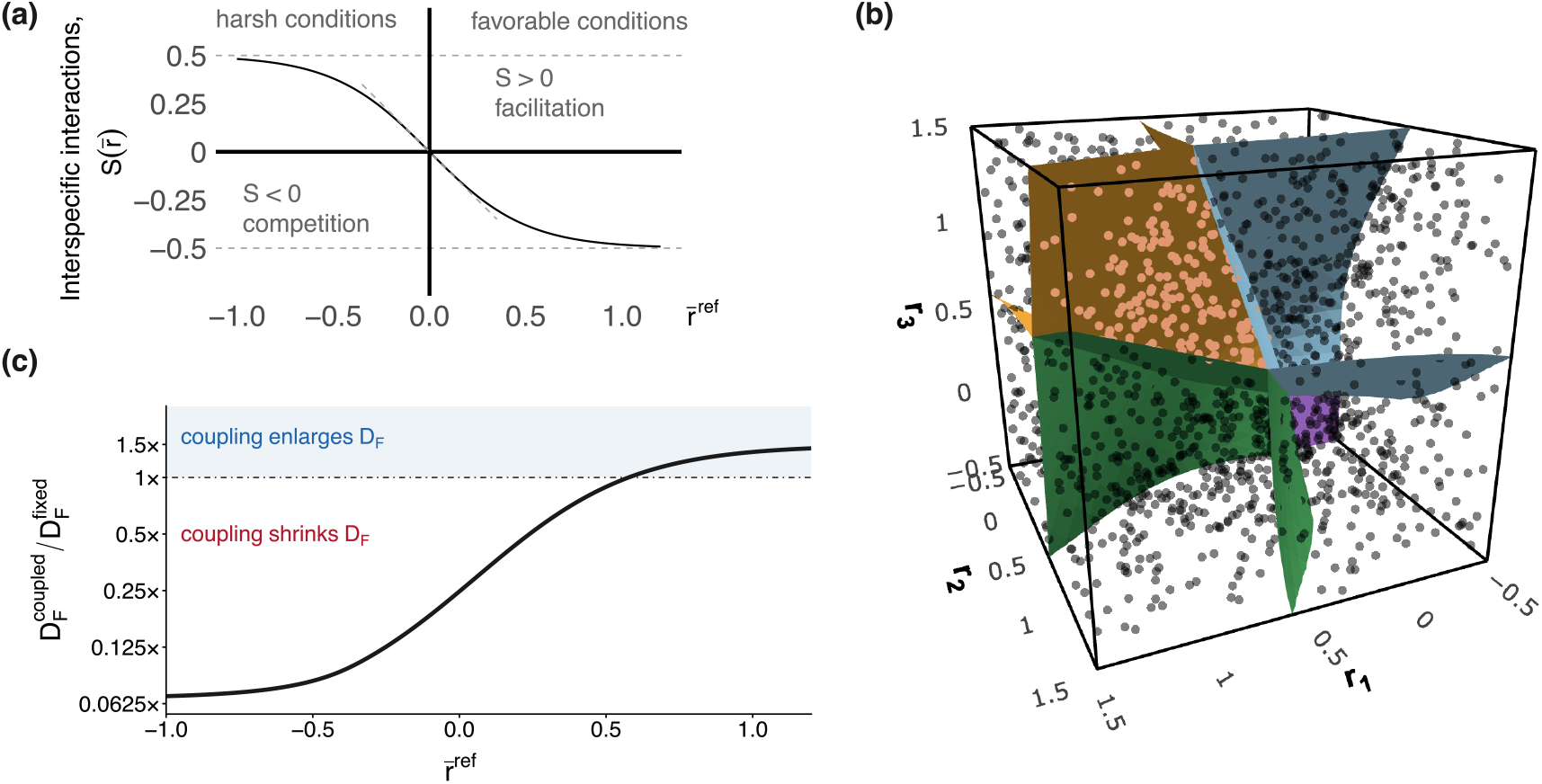
Non-linear interaction coupling via sigmoidal functions. **a)** Sigmoidal function 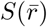 defining the off-diagonal terms of the interaction matrix (i.e., interspecific interactions) as a function of the mean intrinsic growth rate 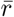. **b)** Intersection of the feasibility domain with the hyperrectangle *r*_*i*_ ∈ [−0.5, 1.5], illustrating volume estimation by Monte Carlo integration (points inside the feasibility domain shown in orange, 2000 in total). **c)** Comparison of the coupled system with the family of uncoupled models it contains. The curve shows the ratio as a fold change on a log scale, and the dash-dotted line at 1 marks equality between the coupled and uncoupled cases, and blue shading indicates the reference conditions for which coupling enlarges the feasibility domain.

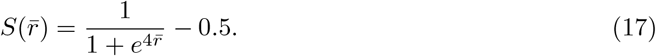

The slope of *S* at its inflection point is −*s*_max_*k/*4, so this choice sets it to −1. Consequently, near 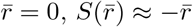, and linear couplings such as the one used for higher-order interactions can be viewed as local approximations of this saturating formulation.

This functional form captures the shift from predominantly positive interactions under stressful abiotic conditions 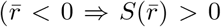, facilitation) to increasingly competitive interactions under high productivity 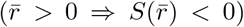, as recently shown for annual plant communities (Buche et al., 2026). The resulting growth-rate-dependent interaction matrix *A*(**r**) for a three-species community is:

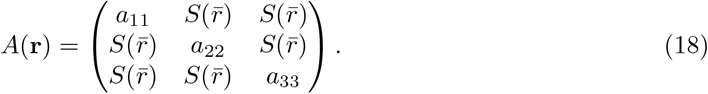

Two simplifying assumptions underlie this formulation for illustration purposes. First, all interspecific interactions are equal and symmetric. Second, they depend on the community mean 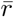 rather than on species-specific growth rates, so that 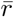 acts as a proxy for the shared environmental conditions or overall productivity experienced by the community. Asymmetric or species-specific couplings can be accommodated by assigning a different sigmoidal function to each species pair.

If intraspecific interactions are fixed or rescaled at −1, 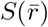 directly measures the strength of interspecific relative to intraspecific interactions (Buche et al., 2025b), therefore isolating the mechanism identified above. That is, how changes in the feasibility domain are driven by how this ratio varies as growth rates increase. Under these assumptions, we estimated the volume of the feasibility domain by Monte Carlo integration within the constraints given by the hyperrectangle *r*_*i*_ ∈ [−0.5, 1.5] Fig. 7b. We analyzed how saturating functional dependencies between growth rates and interactions change the size and shape of the feasibility domain. The coupled system has a single feasibility domain that is 4% of the bounded parameter region, and it lies entirely in the positive orthant (Fig. 7b).

As in the four previous examples, we compare this against the family of uncoupled models, freezing the off-diagonal terms at 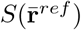 for each reference condition and recomputing the domain over the same constraints. Here, we found that the volume of the uncoupled feasibility domain decreases monotonically as conditions improve and thus interactions turn strongly competitive. The coupled domain is smaller than every member of this family except the most competitive ones, and the two coincide only at 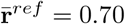, Fig. 7c.

## 4 Main lessons and future directions obtained from coupling ecological strategies

Our main finding is that coupling model parameters through ecological processes fundamentally reshapes the geometry of the feasibility domain and, therefore, rewrites the rules governing the assembly and coexistence of ecological communities across abiotic and biotic barriers. When the interaction matrix becomes a function of intrinsic growth rates, *A*(**r**), the feasibility domain loses the scale invariance that underpins classical structural stability. From a mathematical perspective, this means that it is no longer a polyhedral cone rooted at the origin but a region bounded by curved hypersurfaces. This breaks the standard geometric measure, and therefore the intersection with the unit hypersphere (Saavedra et al., 2017; Song et al., 2020) becomes an arbitrary cross-section rather than a meaningful summary of the feasibility and stability of a community. To solve this limitation, we propose a bounded Monte Carlo estimator, which quantifies structural stability as the proportion of an ecologically realistic hyperrectangle of growth rates that yields feasible and stable equilibria. Critically, our approach can also accommodate couplings that generate forbidden regions inside the feasibility domain, and constraints partitioned into several intervals, both creating holes in the domain (Appendix).

Besides these mathematical considerations, we observe across the first four specific examples of ecological processes that coupling expanded the feasibility domain relative to the uncoupled scenario, regardless of the underlying mechanisms considered. Our different examples should not be seen as separate processes but as processes that represent particular cases of a single facilitation-competition continuum (Buche et al., 2025a), and their common outcome reveals the joint influence of interspecific facilitation and self-limitation on promoting species diversity. At one extreme, the stress-gradient hypothesis implies *a*_*ij*_ *>* 0 under stressful environments, and feasibility extends into regions where *r*_*i*_ *<* 0. This means that species persist despite environments posing strong filtering. At the other extreme, HOIs modify interspecific interactions from competitive *a*_*ij*_ to almost negligible as the initiator species thrives, and so does productivity-mediated niche packing as the community becomes more productive. Both facilitation and competition examples expand the ranges of opportunities for species to coexist because they all increase the ratio |*a*_*ii*_|*/*|*a*_*ij*_| as intrinsic growth rates increase. We believe our results contribute to reducing the long-standing division in the literature that considers facilitation and competition as opposing forces. With our geometric approach where parameters are coupled, both represent alternative routes to the same outcome, and we should not view coexistence as more constrained in communities dominated by facilitation compared to communities in which competition is prevalent and vice versa. Together, these results show that feedbacks between the abiotic and biotic barriers systematically relax the constraints on diversity that uncoupled models predict, and the key question to address in community assembly is not whether a system falls in a given part of the facilitation-competition continuum but rather how assembly changes as species demographic performance changes.

However, when we present the common currency to model all these four ecological processes using the sigmoidal function of changes from positive to negative interactions, we observe the opposite pattern. That is, there is a contraction rather than an expansion of the feasibility domain. Therefore, a fair question to ask is whether there is a way to distinguish expansions from contractions in the opportunities for species to coexist when species’ growth rates and interactions are coupled. The answer to this question is straightforward. The definition of the function for coupling in the *A*(**r**) matrix will determine whether feasibility boundaries bend outward or inward, and this difference basically depends on the sign of the induced change in the relative strength of intra versus interspecific interactions along the direction in which growth rates vary. This can be made explicit by differentiating the equilibrium condition **r** + *A*(**r**)**N**^*^ = 0 with respect to the growth rate of species *i, r*_*i*_, which gives **e**_*i*_ + (*∂A*(**r**)*/∂r*_*i*_) **N**^*^ + *A*(**r**) *∂***N**^*^*/∂r*_*i*_ = 0, and hence:

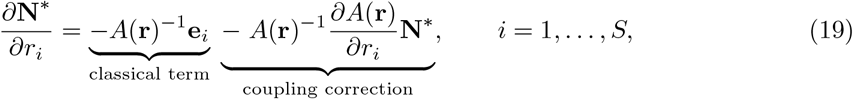

where **e**_*i*_ denotes the *i*-th standard basis vector of ℝ^*S*^, and *∂A*(**r**)*/∂r*_*i*_ is the *S* × *S* matrix whose (*p, q*)-th element is

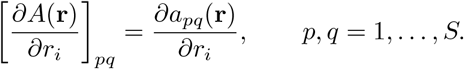

The first term of Eq. (19) recovers the classical, uncoupled sensitivity of equilibrium abundances to demography. The second term is the coupling correction, and its sign relative to the first governs whether the domain expands or contracts. For the first four of our scenarios, the correction increases the strength of self-limitation compared to interspecific competition as growth rates increase. In other words, the matrix of species interactions tends to be diagonally dominant as intrinsic growth rates increase. However, for the sigmoidal function, the correction is the opposite, something that can also likely occur in nature. For instance, if superior competitors take advantage of resources and use them better than their neighbors and grow faster, as in the case of invasive species (Godoy et al., 2014), or conversely, if facilitation collapses at the extreme end of a stress gradient (Michalet et al., 2006), the same formalism predicts a feasibility domain smaller than the uncoupled scenario. Our five scenarios, therefore, illustrate both directions of a general phenomenon that accounts for non-linear changes in the ecological outcomes of coexistence and exclusion.

Taken together, the framework we present here is indeed theoretical, and its natural extension is to expand it to empirical studies. We have shown the ecological implications when demography and interactions are coupled through our toy examples. However, we cannot yet say how often such coupling occurs in nature, how strong it typically is, or which functional form is most ecologically realistic in determining the direction and extent of curvature in the boundaries of structural stability. Answering these questions requires estimating *A*(**r**) rather than *A*. This implies that we should identify environmental or ecological drivers that generate variation in intrinsic growth rates and test whether this variation is accompanied by systematic changes in interaction coefficients. Experimentally, manipulating environmental severity, resource availability, productivity, or the density of several species in diverse communities provides a tractable way to test whether these drivers jointly affect growth rates and interaction coefficients, and whether the resulting dependencies are detectable, consistent, and strong enough to matter. We expect the answer to be context-dependent, and so ecological coupling will either expand or contract the feasibility domain depending on the particularities of the system. This means that the relative prevalence is an open empirical question rather than something theory alone can settle. What our mathematical explorations do establish is that this question cannot be set aside. If parameters are coupled and we analyze communities as though they were not, we are likely reaching wrong conclusions about which communities can assemble and how species coexist. Therefore, ecologists working on community assembly should treat the independence of demographic and interaction parameters as a hypothesis to be tested rather than a modeling convenience to be assumed.

## Acknowledgments

We thank Carlos Gómez-Ambrosi for insightful comments and discussions on the manuscript. Project *PID2024-156228NB-I00* from *Ministerio de Ciencia, Innovación y Universidades* partially supported FST, JAL and JRP.

## Data availability statement

No empirical data has been used for this study. The code for reproducing the figures of this manuscript is available at 10.5281/zenodo.23315875.

## AI Use Statement

ChatGPT (OpenAI) and Claude (Anthropic) was used during manuscript preparation to assist with code development, debugging, and language refinement. All outputs were reviewed and validated by the authors, who are solely responsible for the ideas, analyses, interpretations, and conclusions presented in this manuscript.

