## Appendix for "Coupling demography and interactions reshapes the structural stability of ecological communities"

October 16, 2026

Figures S1–S4 are generated by the R script `Fig_Appendix.R`, hosted in zenodo repository [10.5281/zenodo.23315875](https://zenodo.org/record/23315875).

#### 1 A continuous coupling that opens a hole in the feasibility domain

In the examples of the main text, the coupling functions to illustrate the different mechanisms were selected to be continuous and the resulting feasibility domains have therefore no holes. For examples, the curved boundaries in Fig. 3a (main text) can be written in closed form. For two species, solving $\mathbf{r} + A(\mathbf{r})\mathbf{N}^* = 0$ gives $N_1^* = [a_{12}(\mathbf{r})r_2 - a_{22}(\mathbf{r})r_1] / \det A(\mathbf{r})$ and $N_2^* = [a_{21}(\mathbf{r})r_1 - a_{11}(\mathbf{r})r_2] / \det A(\mathbf{r})$, so species $i$ goes extinct where the numerator of $N_i^*$ vanishes. With the parameterization used in Fig. 3a, ($a_{11} = a_{22} = -1$, $a_{12} = (c - r_1)/2$, $a_{21} = (c - r_2)/2$, $c = 0.99$), $\det A(\mathbf{r}) = 1 - (c - r_1)(c - r_2)/4$ remains positive over the whole bounded region (its minimum, at $r_1 = r_2 = -1$, is $\approx 0.01$). Hence a unique interior equilibrium exists everywhere, and feasibility reduces to $2r_1 + cr_2 - r_1r_2 > 0$ and $2r_2 + cr_1 - r_1r_2 > 0$. The two extinction boundaries are therefore the hyperbolas

$$N_1^* = 0 : \quad r_1 = -\frac{cr_2}{2 - r_2}, \quad N_2^* = 0 : \quad r_2 = -\frac{cr_1}{2 - r_1}. \quad (\text{A1})$$

Close to the origin they reduce to the straight boundaries of the constant matrix $A(\mathbf{0})$, for instance $r_1 \approx -(c/2)r_2$. As the growth rate of the other species increases, the factor $1/(2 - r_j)$ pushes each boundary outward, into the region where the extinct species has a negative intrinsic growth rate. Integrating these two inequalities exactly over the bounded region gives $D_F^{\text{coupled}} = 0.553$, the same value obtained with the Monte Carlo estimator.

##### 1.1 Continuity of the coupling does not rule out holes in the feasibility domain

However, having a continuous feasibility domain is not a consequence of continuous coupling functions. A continuous, and even infinitely differentiable, coupling $A(\mathbf{r})$ can produce a feasibility domain with holes depending on whether abundances become unfeasible.

###### 1.1.1 Continuous coupling function: hump-shaped

Consider two species with an effect of species 2 on species 1 that intensifies within a restricted window of environmental conditions centered at $\mathbf{r}_c$, what we will call a *competition hotspot* when one species (species 2 in this case) reaches its performance optimum:

$$a_{12}(\mathbf{r}) = -a_0 - h \max \left\{ 0, 1 - \frac{\|\mathbf{r} - \mathbf{r}_c\|^2}{\rho^2} \right\}. \quad (\text{A2})$$

This coupling is continuous (Fig. S1a), where the rest of the parameters are: self-regulation $a_{11} = a_{22} = -1$, a constant effect of species 1 on species 2, $a_{21} = -0.2$, $a_0 = 0.2$, $h = 1.5$, $\rho = 0.4$ and $\mathbf{r}_c = (1, 1)$. The coupling is quadratic inside the disk of radius $\rho$ centered at $\mathbf{r}_c$, where competition rises up to $a_{12}(\mathbf{r}_c) = -1.7$, and equals the baseline value $-a_0$ outside it. Ecologically, it describes interspecific competition that peaks at intermediate environmental conditions, as in hump-shaped relationships between productivity and competition intensity, and remains weak elsewhere.

For two species, $N_1^* = (r_1 + a_{12}(\mathbf{r}) r_2) / \det A(\mathbf{r})$ and $N_2^* = (r_2 + a_{21} r_1) / \det A(\mathbf{r})$, with $\det A(\mathbf{r}) = 1 - a_{12}(\mathbf{r}) a_{21} \geq 0.66$ over the bounded region $\mathcal{R} = [-0.5, 2]^2$. A unique interior equilibrium therefore exists everywhere in $\mathcal{R}$ and, since $a_{ii} < 0$ and $\det A(\mathbf{r}) > 0$, it is locally stable whenever it is feasible. Away from the hotspot, feasibility reduces to the classical cone $0.2 r_2 < r_1 < 5 r_2$. Inside the hotspot, species 1 goes extinct on the closed curve

$$r_1 = \left[ a_0 + h \left( 1 - \frac{\|\mathbf{r} - \mathbf{r}_c\|^2}{\rho^2} \right) \right] r_2, \quad (\text{A3})$$

which encloses $\mathbf{r}_c$. The feasibility domain thus contains a hole: a region of growth rates in which species 1 is excluded, completely surrounded by growth rates that allow coexistence (Fig. S2a). Along the diagonal $r_1 = r_2 = r$, for instance, $N_1^* < 0$ exactly when $a_{12}(t, t) < -1$, that is, for $r \in (0.807, 1.193)$ (Figs. S1b and S3). Both an improvement and a deterioration of environmental conditions therefore restore coexistence.

#### 1.1.2 Continuous coupling function: Gaussian-shaped

The hole is not an artifact of the quadratic cap at $\|\mathbf{r} - \mathbf{r}_c\| = \rho$. Replacing it by the smooth Gaussian hotspot

$$a_{12}(\mathbf{r}) = -a_0 - h \exp\left(-\frac{\|\mathbf{r} - \mathbf{r}_c\|^2}{2\sigma^2}\right), \quad \sigma = \rho/2, \quad (\text{A4})$$

gives the same qualitative picture with a slightly smaller hole (Fig. S2b); on the diagonal, species 1 is excluded for $r \in (0.841, 1.159)$. Finally, the bounded Monte Carlo estimator of the main text handles the hole without any modification (Fig. S4). Without the hotspot ($h = 0$), the domain occupies a fraction $\Omega_{\mathcal{R}} = 0.512$ of $\mathcal{R}$. With the quadratic hotspot, the exact value, obtained by integrating the boundary (A3) in polar coordinates around $\mathbf{r}_c$, is $\Omega_{\mathcal{R}} = 0.4736$, and the Monte Carlo estimate with $K = 10^6$ points is $\hat{\Omega}_{\mathcal{R}} = 0.4735 \pm 0.0005$ (one standard error). The difference with respect to the case without hotspot is the area of the hole, 0.240, that is, 3.8% of $\mathcal{R}$.

### 1.2 Figures

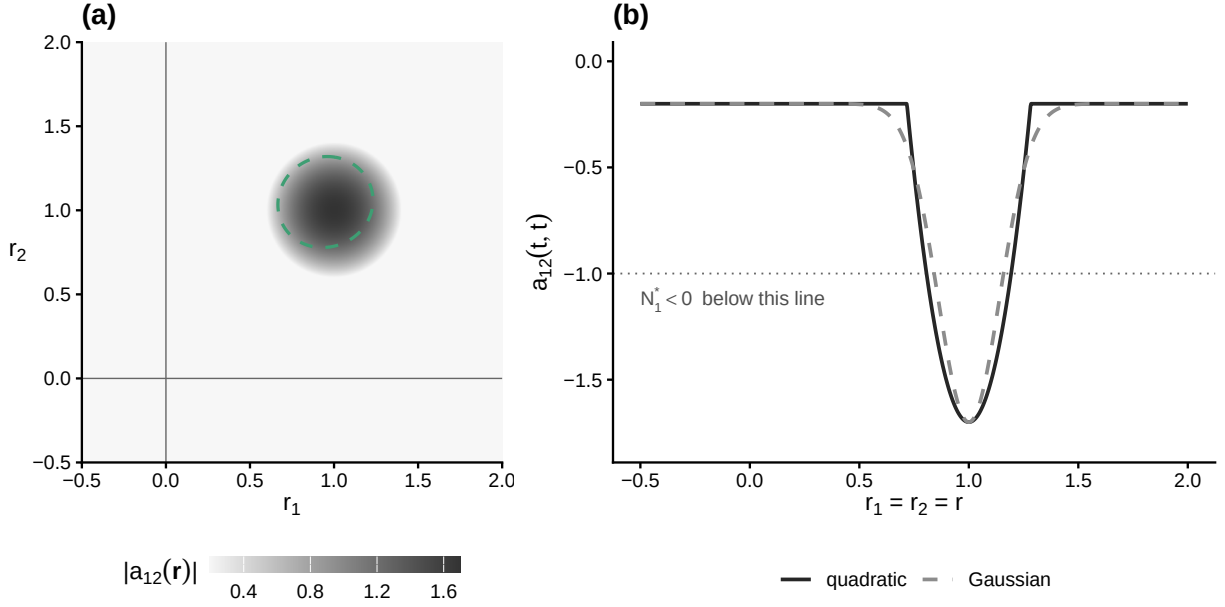

Figure S1: **Competition hotspot.** (a) Strength of the effect of species 2 on species 1, $|a_{12}(\mathbf{r})|$, for the quadratic hotspot of Eq. (A2) over the bounded region $\mathcal{R}$. The dashed curve is the boundary of the hole, Eq. (A3). (b) $a_{12}$ along the diagonal $r_1 = r_2 = r$ for the quadratic (solid) and Gaussian (dashed, Eq. (A4)) hotspots. On the diagonal, $N_1^* < 0$ wherever $a_{12} < -1$ (dotted line).

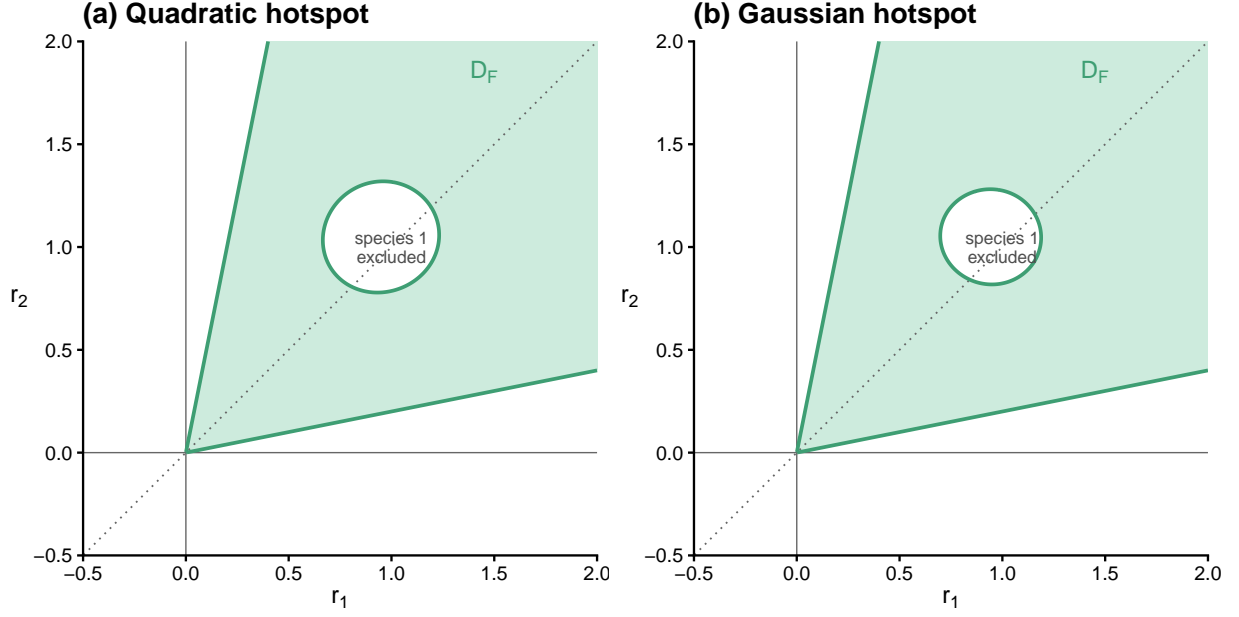

Figure S2: **Feasibility domain with a hole.** Feasibility domain $D_F$ (green) within $\mathcal{R}$ for (a) the quadratic hotspot and (b) the Gaussian hotspot. Green curves are the extinction boundaries. The white region enclosed by the inner curve is the hole: species 1 is excluded there, although every surrounding growth-rate vector allows coexistence. The dotted line is the diagonal $r_1 = r_2 = r$ used in Fig. S3.

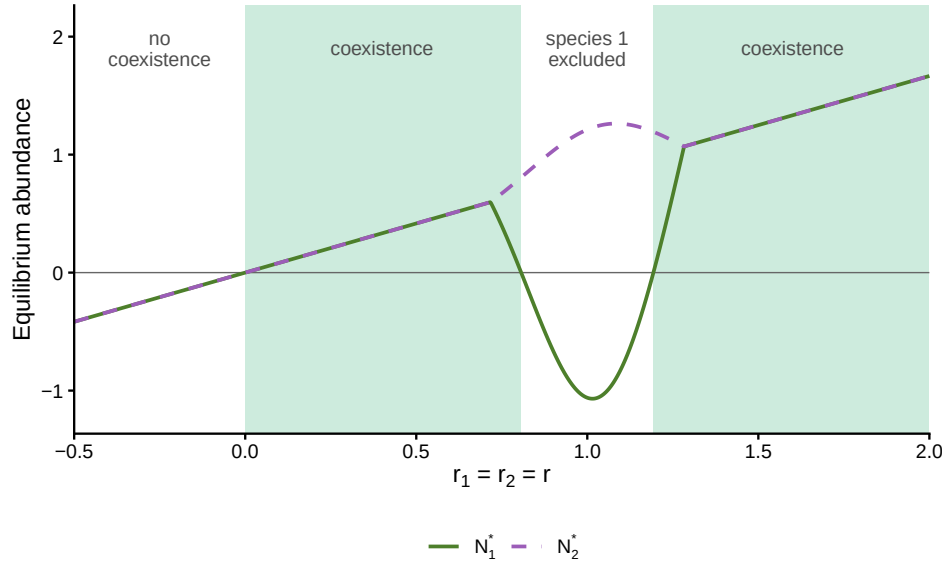

Figure S3: **Loss and recovery of coexistence along an environmental gradient.** Equilibrium abundances along the diagonal $r_1 = r_2 = r$ for the quadratic hotspot. Green bands mark coexistence ($N_1^*, N_2^* > 0$). Outside the hotspot the two species are symmetric and the two curves coincide. Inside it, $N_1^*$ becomes negative for $r \in (0.807, 1.193)$: species 1 is excluded at intermediate conditions, and coexistence is recovered when conditions either improve or deteriorate.

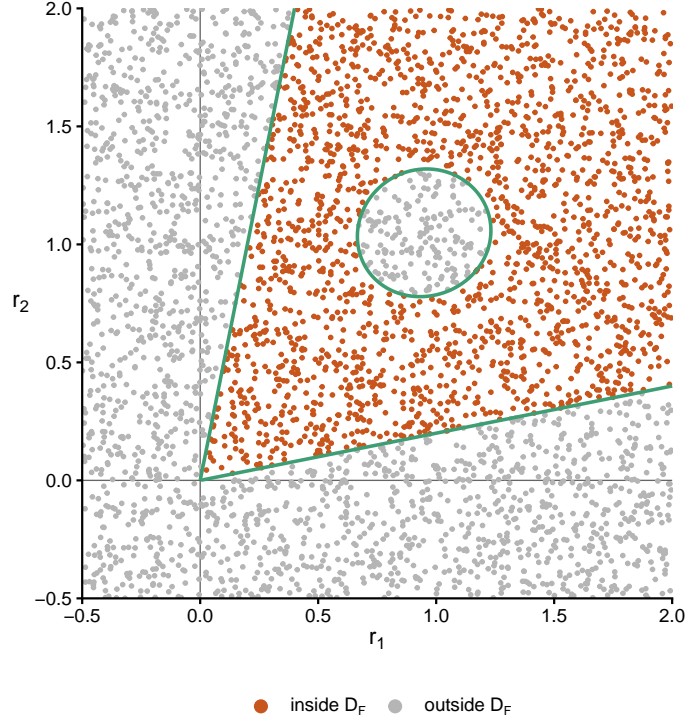

Figure S4: **Monte Carlo estimation in the presence of a hole.** Growth-rate vectors drawn uniformly over $\mathcal{R}$ (4000 points for illustration; the estimate in the text uses $10^6$), coloured by whether they fall inside or outside $D_F$ for the quadratic hotspot. The estimator needs no knowledge of the shape of $D_F$ and simply counts the points in the hole as infeasible.
